# Mirtazapine treatment in a young female mouse model of Rett syndrome identifies time windows for the rescue of early phenotypes during development

**DOI:** 10.1101/2021.12.17.473107

**Authors:** Javier Flores Gutiérrez, Giulia Natali, Jacopo Giorgi, Elvira De Leonibus, Enrico Tongiorgi

## Abstract

Rett Syndrome (RTT) is a rare X-linked neurodevelopmental disorder, mainly caused by mutations in the *MECP2* gene. Reduction in monoamine levels in RTT patients and mouse models suggested the possibility to rescue clinical phenotypes through antidepressants. Accordingly, we tested mirtazapine (MTZ), a noradrenergic and specific-serotonergic tetracyclic antidepressant (NaSSA). In previous studies, we showed high tolerability and significant positive effects of MTZ in male *Mecp2*^1m1.1Bird^-knock-out mice, adult female *Mecp2*^tm1.1Bird^-heterozygous (*Mecp2*^+/-^) mice, and adult female RTT patients. However, it remained to explore MTZ efficacy in female *Mecp2*^+/-^ mice at young ages. As RTT-like phenotypes in young *Mecp2*^+/-^ mice have been less investigated, we carried out a behavioural characterization to analyze *Mecp2*^+/-^ mice in “early adolescence” (6 weeks) and “late adolescence/young adulthood” (11 weeks) and identified several progressive phenotypes. Then, we evaluated the effects of either a 15- or a 30-day MTZ treatment on body weight and impaired motor behaviours in 11-week-old *Mecp2*^+/-^ mice. Finally, since defective cortical development is a hallmark of RTT, we performed a histological study on the maturation of perineuronal nets (PNNs) and parvalbuminergic (PV) neurons in the primary motor cortex. The 30-day MTZ treatment was more effective than the shorter 15-day treatment, leading to the significant rescue of body weight, hindlimb clasping and motor learning in the accelerating rotarod test. Behavioral improvement was associated with normalized PV immunoreactivity levels and PNN thickness. These results support the use of MTZ as a new potential treatment for adolescent girls affected by RTT and suggest a possible mechanism of action.

## INTRODUCTION

Rett Syndrome (RTT; OMIM 312750) is a rare, severe X-linked progressive neurodevelopmental disorder *(1, 2)*. With an incidence of 1/10,000 newborn females, RTT is the second leading cause of intellectual disability of genetic basis in girls *(3)*. In most cases of its classic form, RTT is caused by a mutation of the *MECP2* gene (methyl-CpG binding protein 2, HGNC:6990) *(4)* and more than 100 mutations have been described with extensive clinical heterogeneity *(5–7)*. MeCP2 protein is a transcription factor that acts as a context-dependent global organizer of chromatin architecture, activating or down-regulating the transcription of numerous genes, but in recent times additional functions have been attributed to this protein in the regulation of splicing, miRNA processing, and protein synthesis *(8, 9)*. Despite high genetic variability, RTT patients undergo similar clinical trajectories. After 6-24 months of apparently normal development, RTT patients lose previously acquired skills such as the purposeful use of hands and verbal language, manifesting typical stereotypic hand movements and autistic traits *(5)*. Then, slower growth of the brain occurs, resulting in microcephaly, loss of walking, tremor, and other movement disorders (e.g. dystonia and ataxia) *(10)*. At this phase, epilepsy frequently appears, ranging from easily controlled to intractable seizures *(11)*. Within adolescence, social interactions tend to improve, but somatic growth restriction and scoliosis usually become more evident. Constipation and breathing abnormalities are also present frequently at this phase. In adulthood, motor deterioration may increase, often leading to parkinsonism *(12)*. Various clinical trials have been carried in RTT patients without significant benefit *(13)*. Thus, at present, there is still no cure for RTT.

The observed reduction in monoamine levels (NE, 5-HT, DA) in RTT patients and mouse models suggested the possibility to rescue the clinical phenotype through antidepressants *(14)*. Accordingly, in a previous study, we tested mirtazapine (MTZ), a noradrenergic and specific-serotonergic tetracyclic antidepressant (NaSSA) with high tolerability and limited side effects *(15–19)*. We found that 2-week treatment of one-month-old male *Mecp2*-null mice with 10 or 50 mg/kg MTZ rescued both heart and breath rate deficits without cardiovascular side effects *(20)*. In addition, MTZ re-established the normal glutamatergic and GABAergic transmission in both cortex and brainstem, and rescued microcephaly by restoring the fine neuronal morphology and the somatosensory cortex thickness *(20)*. However, this starting study was performed in male *Mecp2*-null mice, which are an imperfect model of the pathology because they present an early severe phenotype and die very young. In addition, they are not a genetic mosaicism as female RTT patients and female mice. Therefore, we performed a study in adult heterozygous female *Mecp2*^tm1.1Bird^ (*Mecp2*^+/-^) mice and adult female RTT patients, and we demonstrated that MTZ has protective effects against motor deterioration and other behavioural features since the progression of the symptoms was halted, or in some cases even reverted *(21)*. However, considering that adults have reduced plasticity of the nervous system compared to younger subjects, it remained to verify the possibility that MTZ might have higher beneficial effects in female *Mecp2*^+/-^ mice at earlier ages.

Given the substantial gap in the knowledge of early behavioural phenotypes of female *Mecp2*^+/-^ mice, we used a battery of tests to analyze the Bird RTT murine model *(22–27)* at two early stages of development, i.e. in “early adolescence” (6 weeks, soon after mice completed their motor development) and in “late adolescence/young adulthood” (11 weeks of age, after sexual maturation) *(28)*. Then, we evaluated the efficacy of either a 15-day or a 30-day daily treatment with 10 mg/kg MTZ in 11-week-old female mice, using tests for general motor coordination, motor learning and progression of hindlimb clasping, which resembles the sensorimotor alterations observed in RTT patients. In addition, we performed a histological study on perineuronal nets (PNNs) in the primary motor cortex. The results obtained support MTZ as a new potential treatment for children and adolescent girls affected by RTT.

## MATERIALS AND METHODS

### Animals

Animals were treated according to the institutional guidelines, in compliance with the European Community Council Directive 2010/63/UE for care and use of experimental animals. Authorization for animal experimentation was obtained from the Italian Ministry of Health (Nr. 124/2018-PR), in compliance with the Italian law D. Lgs.116/92 and the L. 96/2013, art. 13. All efforts were made to minimize animal suffering and to reduce the number of animals used. For animal production, *Mecp2* heterozygous (HET) females (*Mecp2*^+/-^, B6.129P2(C)-*Mecp2*^tm1.1Bird/J^, Stock No: 003890. The Jackson Laboratory, USA) *(22)* were bred with wild-type C57/BL6J male mice (The Jackson Laboratory, USA). We used female *Mecp2*^*+/-*^ mice at 6 and 11 weeks of age to characterize the phenotypes at different developmental stages. After weaning, wild-type (WT) and heterozygous (HET) mice were housed in groups (littermates together) in ventilated cages under 12h light/dark cycle with food and water ad libitum. No environmental enrichment elements were added to the cages. All experiments were performed blindly to the genotype and treatment of animals, and all control animals were WT age-matched littermates of HET mice. Finally, mice were assigned to groups according to the rules indicated by Landis et al. *(29)*.

### Mice genotyping

Biopsies from ear punches were incubated with 250 µL of DNA extraction buffer (TRIS 10 mM pH 7.5, EDTA 5 mM pH 8, SDS 0.2%, NaCl 10 mM, proteinase K 0.5 mg/mL) and left overnight at 55°C. The day after, samples were centrifuged (12000 rpm, 20 min, RT), then 100 µL of the supernatant were mixed with isopropanol (1:1), and precipitated DNA was centrifuged again (12000 rpm, 30 min, RT). The supernatant was then discarded and three washes with cold 70% ethanol with subsequent centrifugations (12000 rpm, 5 min, RT) were realized. Once ethanol had evaporated, DNA pellets were homogeneously dissolved in milliQ water. Genotypes were assessed by PCR on genomic DNA extracted from ear-clips biopsies. PCR reactions were performed using specific primers (forward common primer oIMR1436 5′-GGT AAA GAC CCA TGT GAC CC-3′, reverse mutant primer oIMR1437 5′-TCC ACC TAG CCT GCC TGT AC-3′) with 1U GoTaq polymerase (Promega, Madison, USA), 1X green GoTaq buffer, 0.2 mM dNTPs each, 2.5 mM MgCl_2_, 0.5 µM of each primer and 10 ng/µL of genomic DNA, as follows: 95°C, 3’ > 30 cycles: 95°C, 20’’; 58°C, 20’’; 72°C, 20’’ > 72°C, 2’. This PCR generates a 400-bp product for the WT allele and an additional 416-bp product for heterozygous mice *(20)*.

### Mice treatment

Beginning from 9 weeks of age, HET mice and WT littermates were i.p. injected with vehicle (VEH = 0.9% aqueous solution of NaCl and 5% ethanol) or MTZ (10 mg/kg, ab120068, Abcam, Cambridge, UK). The 10 mg/kg daily dosage in mice is equivalent to 50 mg/day in humans, which is the maximum dose used in patients *(15, 30)* In addition, this dosage has shown no toxicity in adult HET mice *(21)*. To calculate it, we followed a dose by factor method modified from Nair and Jacob *(30)*. The next formula was used:

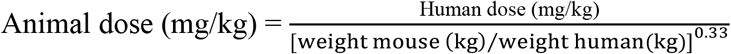

We used standard weights described by authors *(30)* (mouse = 0.02 kg; human: 60 kg), obtaining a dose of 11.7 mg/kg, which we rounded to 10 mg/kg. Mice were treated every day at 10-11 a.m., for either 15 or 30 days. As general phenotypic scoring *(21)* did not show significant differences between mice at the beginning of the experiment, we just divided randomly WT and HET mice into two groups, VEH- and MTZ-treated, thus creating our 4 experimental groups (WT-VEH, WT-MTZ, HET-VEH and HET-MTZ).

### Animal behaviour

All tests were performed the day after the last i.p. injection had been applied (off-drug). A timeline including every used test is shown in Suppl. Fig. 1. The order of the tests we followed aimed to minimize interferences among them: elevated plus-maze (EPM), accelerating rotarod (for three consecutive days), four-different objects task (4-DOT) and rod walk test. Every test was performed on a different day. In addition, body weight and hindlimb clasping were weekly assessed, starting the first day of treatment. Finally, adhesive patch removal task was performed only for behavioural characterization of untreated animals.

### Hindlimb clasping

This abnormal movement is a marker of disease progression in several mouse models of neurological diseases, including RTT *(27)*. To assess it, we grasped the tail near its base and lift the mouse clear of all surrounding objects, and then observed the hindlimb position for 10 seconds. We assigned a score of 0 when hindlimbs were consistently splayed outward, away from the abdomen. A score of 1 was assigned when one hindlimb was retracted towards the abdomen for more than 50% of the time suspended. When both hindlimbs were either partially or entirely retracted toward the abdomen for more than 50% of the time suspended, we assigned a score of 2.

### Elevated-plus maze (EPM)

Each mouse was individually placed in the centre area of a black Plexiglass elevated-plus maze for a 5-min test session *(21, 26)*. Mice movements were recorded and later analysed with ANY-maze software (Stoelting, New Jersey, USA) to automatically measure entries and time spent in the open and in the closed arms.

### Accelerating rotarod test

This test evaluates general motor coordination and motor learning in mice, through three trials per day (separated by a one-hour intertrial interval), which were repeated over three consecutive days. In each trial, the mouse was placed on the rotarod apparatus (Ugo Basile, Varese, Italy) and then the rotation started at 5 revolutions per minute (rpm). After the mouse was successfully walking, the rotarod was accelerated from 5 to 40 rpm over 5 minutes. The latency to fall from the rotating beam to the flange floor of the apparatus was recorded for each animal. The trial was terminated in cases where the mouse clutched the beam without walking, for three consecutive turns, or a maximum of 300 s had elapsed *(21, 26)*

### Four-different objects task (4-DOT)

We slightly modified a protocol published by Sannino et al. *(31)*, in which three different phases of the task are described. Starting three days before the test, mice were daily habituated to new objects (different from experimental ones) and to individual cages. On the day of the test, each mouse was first left in the centre of an open field arena (35 × 47 × 60 cm) and allowed to freely explore it for 10 minutes (habituation phase, T1). Then the mouse was put back in its individual cage and left there for 1 min. In the meanwhile, four different objects were placed symmetrically in the open field arena. Then the mouse was put again into the open field arena and left to explore objects for 5 min (familiarization phase, T2). After 10 min, the mouse was put back in the isolation cage for 1 min and, in the meanwhile, all objects were put away. Three of them were substituted by identical copies and just one was substituted by a completely different object (the novel object). Each mouse was finally left to explore this new set of objects for 5 min (test phase, T3). All phases were completely video-registered and then analysed with ANY-maze software (Ugo Basile Instruments). The exploration (sniffing) time for each object was manually measured in the whole T2 and in the whole T3. We then calculated the re-exploration index for each object and each mouse, which is the result of subtracting the exploration time in T2 to the exploration time in T3. In this way, a negative re-exploration index indicates that the mouse recognizes the specific object as already known. On the contrary, a positive re-exploration index indicates a higher interest for the object, which in WT mice corresponds to the normal exploration of a novel object.

### Rod walk test

Following a modified protocol based on *(32)* and already tested in adult *Mecp2* HET female mice *(21)*, we positioned the mouse at one edge of a 60-centimeter wood dowel (standing 50 cm above the floor), where a repulsing stimulus (strong light) was present. On the other side of the dowel, we positioned an attractive stimulus (dark cage with nesting material from the mouse’s home cage). Transition time was measured in two consecutive trials, performed with striped dowels of 10 mm in diameter.

### Adhesive patch removal task

Following a modified protocol of *(33)*, the mouse was allowed to get used to the experimental cage before the trial for one minute. After that, it was immobilized by gripping the scruff of its neck and then one squared adhesive tape strip (0,5 × 0,5 cm) was stuck on each forepaw. The animal was then put back into the experimental cage and thus the time to remove both adhesive patches was measured, (starting when it begins to try to do it).

### Tissue preparation and immunofluorescence procedure

Brains were dissected from mice immediately after the last behavioural test, fixed in PFA and then cryoprotected by immersion in 20% sucrose. They were kept at 4 °C until cutting at the cryostat (CM3050S; Leica Wetzlar, Germany). We produced 20-μm thick coronal sections of the primary motor cortex (M1, approximately from Bregma +1.78 to Bregma +0,62) *(34)* and then collected free-floating in PBS. For each experiment, brains from all experimental groups were processed in parallel. Brain slices were permeabilized with 1% Triton X-100 (1 h, RT) and then incubated in 0,1% Triton X-100 and 1% bovine serum albumin (BSA, Sigma-Aldrich, A7906) for 1,5 h at RT. After that, sections were incubated overnight at 4 °C with primary antibody solution. We use a combination of antibodies (rabbit anti-parvalbumin, 1:5000, PV235, Swant + biotin-conjugated lectin from WFA, 1:250, L-1516, Sigma-Aldrich). On the day after, slices were washed with PBS and incubated in donkey anti-rabbit IgG secondary antibody conjugated with Alexa Fluor 568 (1:250, A10042, Invitrogen) for 1,5 h at RT. Stained sections were rinsed three times (5 min each) in PBS and incubated in 1:1000 Hoechst solution (33342, Sigma) for 20 minutes at RT. Finally, sections were fast washed in PBS and milliQ water and mounted on 76 × 26 mm gelatine-coated microscope slides (Thermo Fisher Scientific), air-dried, and cover-slipped with Mowiol (Merk Life Science). They were finally stored covered from light at 4 °C until microscopy analysis the day after.

### Acquisition of images for histological analyses

By using a Nikon Eclipse TE2000-U confocal microscope equipped with EZ-C1 software (v3.91, Nikon), we acquired images of the primary motor cortex (M1) stained with anti-PV antibodies and lectin from WFA. We used a 20X objective to create 500-μm width digital boxes, spanning from cortical layer II to cortical layer V-VI, and acquiring 1024 × 1024-pixel images. In the Z-axis, we always used a step size of 3,5 μm. In addition, laser intensity and gain of the photomultiplier were set on the WT-VEH (control) and kept for each experiment, to be able to further compare fluorescence intensity among groups.

### Analysis of PV+ cells

By using FIJI software *(35)*, somata of PV+ cells were manually outlined, and then the mean pixel intensity of the PV channel was automatically measured all along the z-stack, together with the average soma area of the slice. Confocal z-stacks were then transformed into 2D pictures by using the ‘maximum projection’ option of the ‘z-project’ menu. Cell counting was manually performed on these pictures by using the ‘cell counter’ plugin. This procedure also allowed us to obtain single × and Y coordinates of the centre of every PV+ cell and to further calculate the neuronal inter-distance.

### Analysis of PNN thickness

For this procedure, 2D images of the PNN and PV channels were used. By using FIJI software *(35)*, we first subtracted the PV channel to the PNN one, thus resulting in a less intense PNN image. Then 3-pixel thick lines transversally crossing PNN were drawn and their correspondent densitometry plots were automatically produced. On each plot, a threshold of 10% of the maximum intensity value was used as a cut-off to measure the thickness of each side of the PNN. For each PNN, thickness values of both sides were averaged.

### Experimental design and statistical analysis

All behavioural experiments were done blindly to the experimenter, and all experimental groups were tested in the same session in a randomized order. Number of neurons, slices and mice evaluated through immunohistological techniques are reported in the specific section. Data analysis and data graphics were performed with GraphPad Prism 8.0 software (GraphPad, La Jolla, California, USA). When appliable, we used the Shapiro-Wilk test to define data as parametric or non-parametric. To test differences between the two groups, either Student’s t-test or Mann-Whitney test was used as appropriate. One-way ANOVA (with the Dunnett’s posthoc test) or Kruskal-Wallis test (with the corrected Dunn’s posthoc multiple comparisons test) were used for comparing multiple groups when just one factor was considered. The combined effect of two independent variables was tested using a standard two-way ANOVA (followed by Tukey’s multiple comparisons test). Just for the analysis of 4-DOT results, we used a two-way ANOVA with repeated measures to compare the re-exploration index, followed by Dunnett’s multiple comparison test. To analyze the three factors of the Accelerating rotarod test in treated mice (genotype × treatment × day) we used a three-way ANOVA with repeated measures followed by Tukey’s multiple comparisons test. All outliers were detected using Grubb’s test. All details on statistical analyses were collected into Suppl. Fig. 2.

## RESULTS

### *Mecp2*^+/-^ mice show behavioural phenotypes at young ages

The RTT mouse model developed by Guy et al. in 2001 *(22)* has been largely used and many different behavioural alterations have been identified *(36)*. However, most of these studies have been carried out on young male mice or adult females, while the phenotype in young female mice has been comparatively less investigated *(26, 37)*. Thus, before testing MTZ treatment in heterozygous (HET) *Mecp2*^***+/-***^ young female mice, we performed a behavioural characterization to identify consistent phenotypes at 6 and 11 weeks of age, which correspond to “early adolescence” (after completing motor development) and “late adolescence/young adulthood” (after sexual maturation), respectively *(28)*. We found that body weight was already significantly increased in HET mice as soon as 6 weeks of age (Fig. 1A). A similar result was observed also in 11-week-old HET mice. Hindlimb clasping was present in most HET mice, showing a statistically significant phenotype at both ages (Fig. 1B). Regarding sensorimotor phenotypes, the horizontal bars test did not show any difference between WT and HET mice at 11 weeks of age (Fig. 1C), unlike what was previously observed in adult HET mice *(21)*, and therefore we did not perform this test at 6 weeks of age. We assessed basal motor performance and motor learning using the accelerating rotarod and the rod walk tests (Fig. 1D-E), and the fine motor skills through the adhesive patch removal task *(38)* (Fig. 1F) at both ages. Motor performance was impaired already at 6 weeks of age as evidenced by reduced transition time in the Rod walk (Fig. 1E), while motor impairment was statistically significant at the accelerating rotarod only at 11 weeks (Fig. 1D). At 6 weeks of age, HET animals show preserved motor learning capacity, as evidenced by a day-dependent increase in the latency to fall in the rotarod, while at 11 weeks their motor learning is fully impaired, as evolution of their performance at the accelerating rotarod demonstrates (Fig. 1D). In the adhesive patch removal task, which evaluates fine motor skills based on the integration of somatosensory inputs, HET mice showed to be impaired only at 11 weeks (Fig. 1F). The aberrant preference for the open arms, previously observed in adult HET mice when tested at the elevated plus maze *(26)* and demonstrated to be due to a heightened whiskers sensitivity *(21)*, was confirmed in young mice at both 6 and 11 weeks of age (Fig. 1G). Finally, we tested young HET mice at the four-different objects task (4-DOT) *(31)*, to evaluate short-term object memory, and observed a clear deficit in the novel object discrimination during the test phase at 11 weeks of age only; in fact, both WT and HET mice were able to distinguish the novel object from the familiar ones at 6 weeks of age (Fig. 1H).

**Figure 1.**
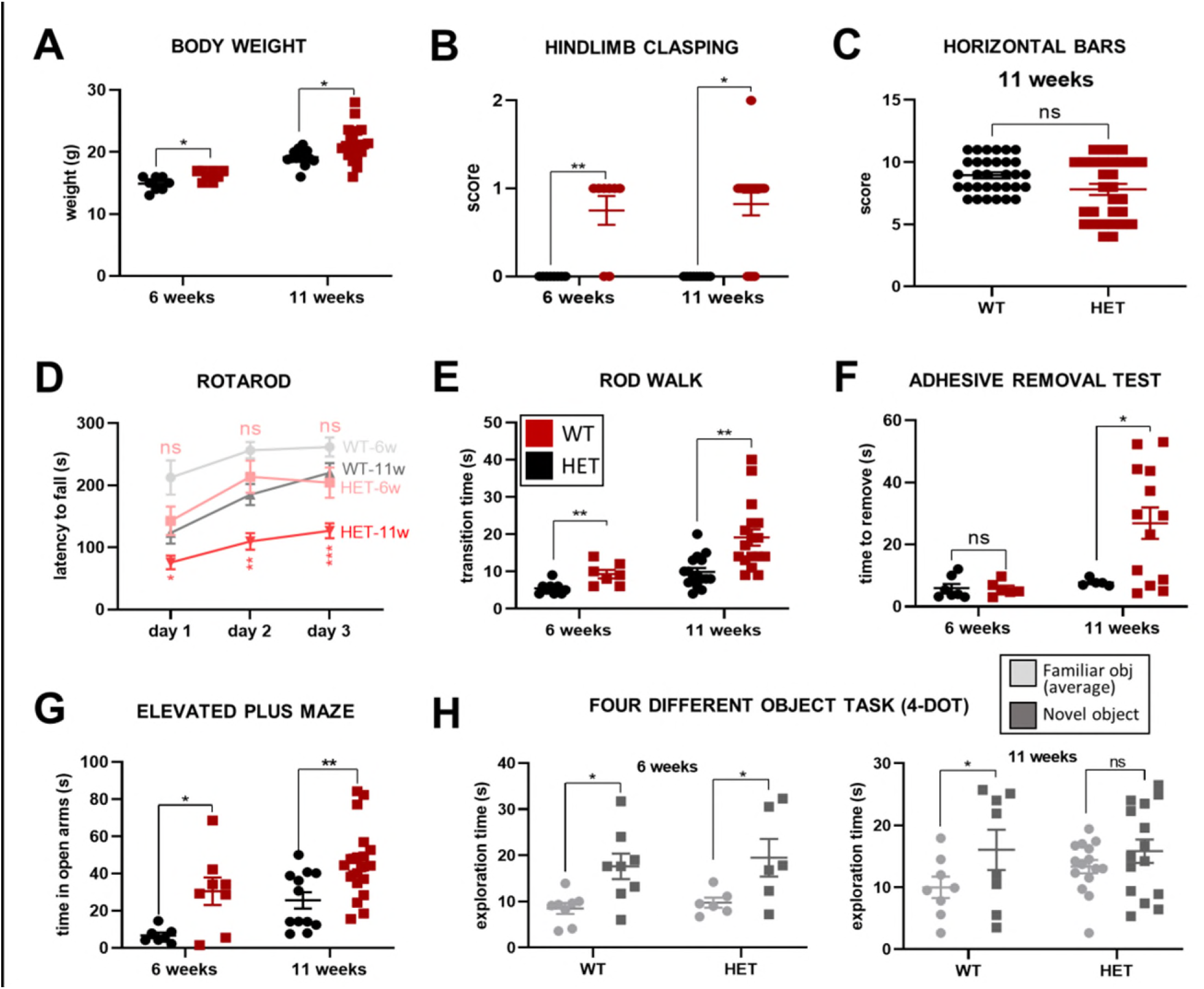
Behavioural phenotypes in *Mecp2*^tm1.1Bird^ female mice at 5 and 11 weeks of age. **(A)** Body weight of mice is shown. **(B)** Hindlimb clasping was evaluated following Guy et al. (2007): 0, absent; 1, mild; 2, strong. **(C-F)** Motor tests. Horizontal bars test was not performed at 5 weeks of age because of absence of phenotype at 11 weeks. In the Accelerating rotarod test, statistical significance was evaluated only for differences between genotypes at the same age. **(G)** Time spent in open arms at the Elevated plus maze (EPM) is shown. **(H)** Time spent exploring familiar and novel objects during the T3 of the Four-different objects task (4-DOT) is shown. Statistical analyses showed that, at 6 weeks, both WT and HET mice were able to recognize the novel object, while at 11 weeks only WT mice were able to do it, this showing a significant phenotype at the older age. All data are expressed as mean ± SEM and all individual values are showed, n = 8-17 mice per each group. According to results of Saphiro-Wilk test, we performed either t-test or Mann-Whitney test. For the Rotarod a standard two-way ANOVA followed by Tukey’s multiple comparison tests was performed, while for the 4-DOT analysis, a two-way ANOVA with repeated measures was used to compare the exploration time of objects, followed by Tukey’s multiple comparison test. ns: p> 0.05, *: p ≤ 0.05, **: p ≤ 0.01, ***: p ≤ 0.001.

In summary, our results identified an optimal time window to test MTZ at 11 weeks of age, when consistent sensorimotor phenotypes are present, but plasticity processes of the nervous system are still significantly active compared with adults. In these pharmacological experiments we have included only those tests that were sensitive to early alteration (i.e. with onset at 6 weeks) in HET mice, as the aim of the study was to test whether MTZ treatment could rescue and not prevent the onset of the behavioural deficits. Accordingly, evaluation of the efficacy of the pharmacological treatment included assessment of body weight, hindlimb clasping, motor learning on rod walking test and accelerating rotarod, and somatosensory sensitivity on the elevated plus maze.

### Body weight is reduced by MTZ treatment

To identify a proper time window for a reliable evaluation of the efficacy of an early treatment, we tested MTZ at 10 mg/kg (the equivalent to the maximum dose in humans) through daily injections for either 15 or 30 days in HET mice, starting treatments at 9 weeks (Suppl. Fig. 1). Body weight was assessed weekly, taking the last measurement the day after the last injection. While we did not observe any statistically significant effect of MTZ treatments after 15 days of treatment (Fig. 2A), administration of MTZ for 30 days led to a statistically significant reduction of the body weight at the end of the treatment (Fig. 2B).

**Figure 2.**
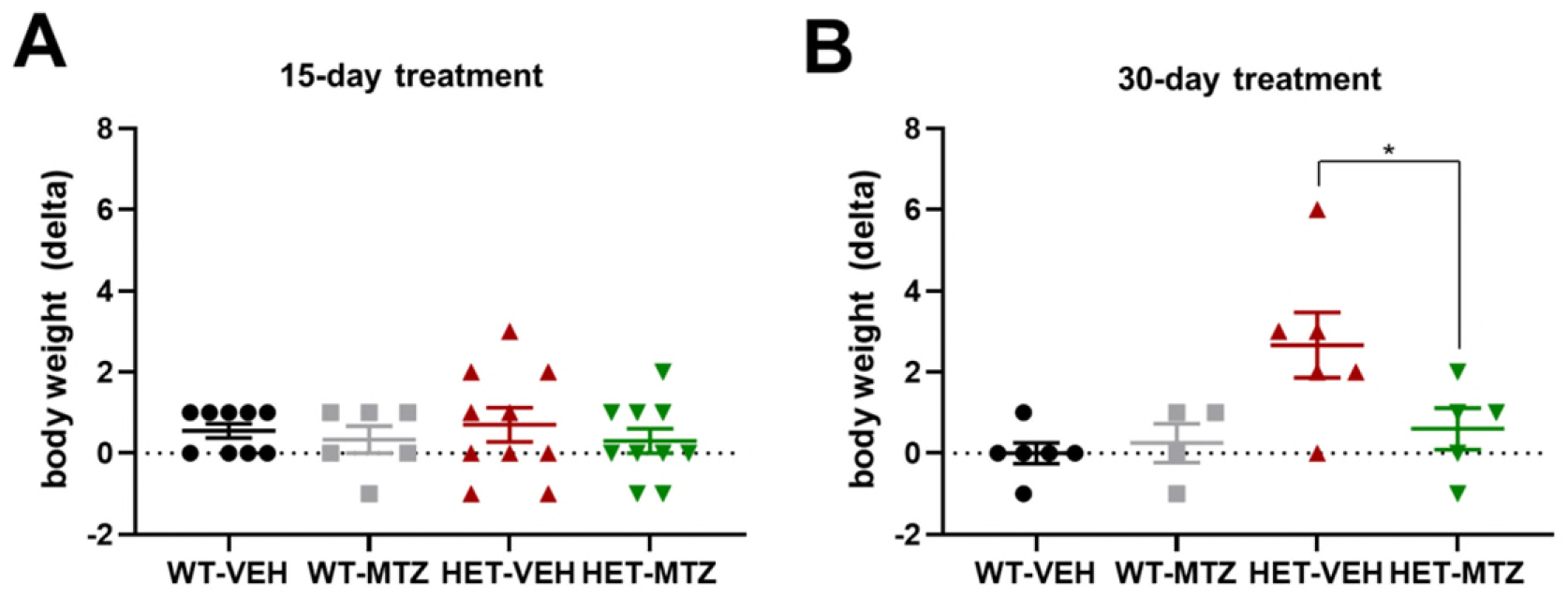
Body weight of 11-week-old *Mecp2*^tm1.1Bird^ female mice after 15 and 30-day treatments with MTZ. In both cases, delta variations were calculated as the difference between the body weight measured at the end and at the beginning of the treatment. All data are expressed as mean ± SEM and all individual values are showed, n = 9-10 mice/group for 15-day analysis and 5-6 mice/group for 30-day analysis. A standard two-way ANOVA followed by Tukey’s multiple comparisons test was performed. ns: p> 0.05 (not indicated), *: p ≤ 0.05, **: p ≤ 0.01, ***: p ≤ 0.001.

### A 30-day MTZ treatment slows down the progression of hindlimb clasping

Together with the body weight, the hindlimb clasping was evaluated weekly by using a scoring system from 0 to 2 (see Methods). When considering the evaluation of this sign only on the last day of treatment, we found that in both groups the scoring of HET was significantly higher (i.e. worst symptom) than that of WT mice (Fig. 3A-B). In addition, after 30 days of treatment with MTZ we found a significant improvement of hindlimb clasping (Fig. 3B), which included 3 out 5 MTZ-treated mice that completely normalized the phenotype. In addition, considering the difference between the score for the hindlimb clasping assigned to the mouse at the last evaluation and the score assigned at the first one (delta values), we confirmed these results, as the phenotype was observed at both ages but only after the 30-day MTZ treatment the delta score in HET mice was significantly lowered (Fig. 3C-D). Taken together, these findings indicate that MTZ is able to slow down the progression of this sign.

**Figure 3.**
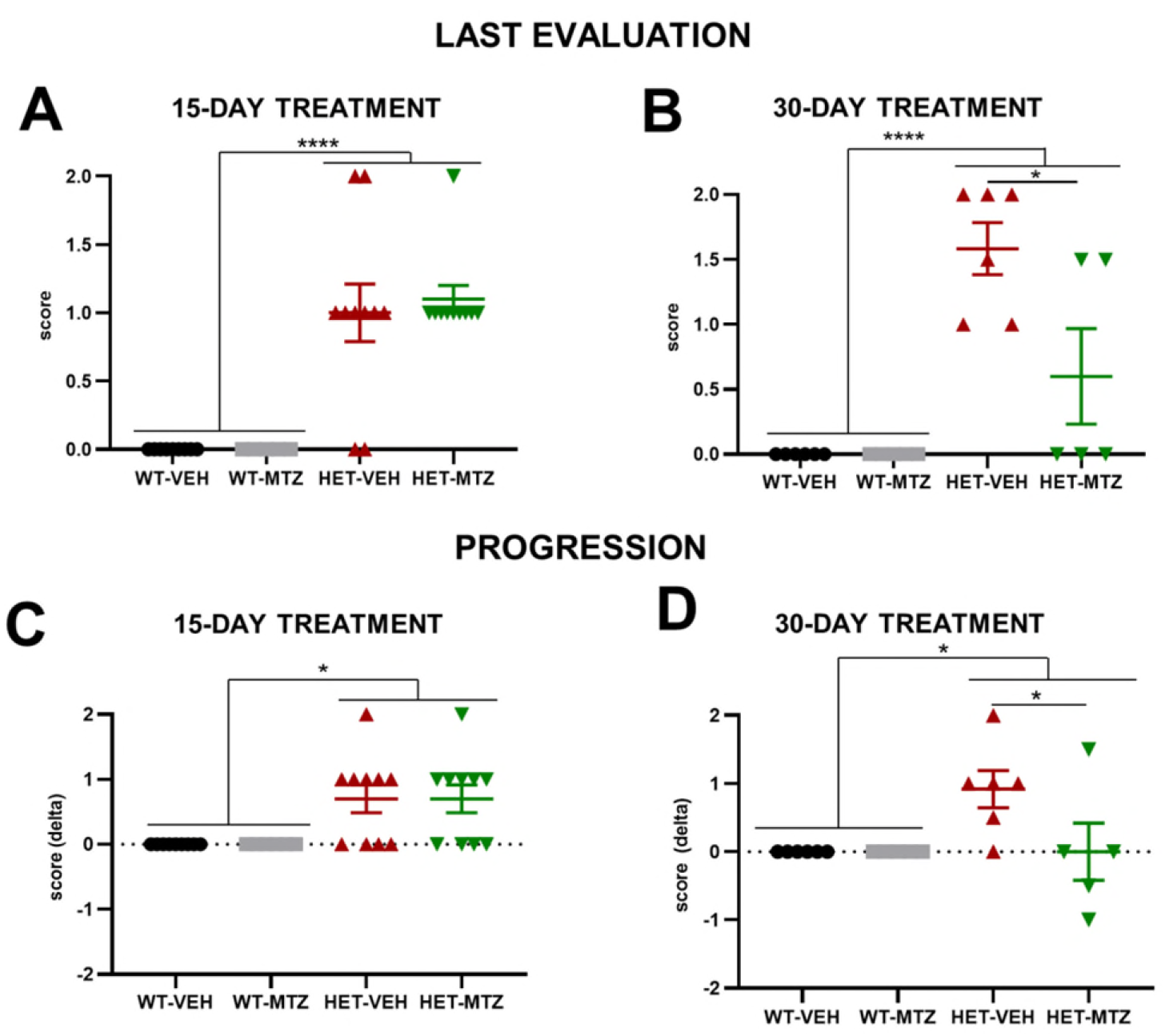
Hindlimb clasping in 11-week-old *Mecp2*^tm1.1Bird^ female mice after a 15 and 30-day treatment with MTZ. **(A-B)** The hindlimb clasping evaluation made after the last injection is shown. **(C-D)** Progression of the hindlimb clasping is shown as delta values. All data are expressed as mean ± SEM and all individual values are showed, n = 9-10 mice/group for 15-day analysis and 5-6 mice/group for 30-day analysis. A standard two-way ANOVA followed by Tukey’s multiple comparisons test was performed. ns: p> 0.05 (not indicated), *: p ≤ 0.05, **: p ≤ 0.01, ***: p ≤ 0.001.

### A 30-day MTZ treatment rescues motor learning

The general motor performance of HET mice was evaluated through two different tests: the rod walk and the accelerating rotarod. In the rod walk, we confirmed the phenotype observed in untreated HET mice (Fig. 1E), as general motor coordination was showed to be impaired in HET-VEH mice, but no effect of MTZ was observed, nor after 15, neither after 30 days of treatment (Fig. 4A-B). In the accelerating rotarod we measured both daily motor performance and 3-day motor learning, this latter is represented by the delta value of the latency to fall (e.g., difference between the value of this parameter on the last day of the test and the value on the first day). We confirmed the deficit of the general motor performance in HET mice, as the latency to fall was significantly shorter compared to WT on the second and the third day of the test (Fig. 4C-D). Whereas the 15-day treatment showed no effect (Fig. 4C), this motor deficit was rescued in HET mice treated with MTZ for 30 days, as they showed a daily performance comparable to that of WT mice (Fig. 4D). In further support of these results, motor learning observed at the accelerating rotarod was rescued by the 30-day MTZ treatment, while the 15-day one showed no effect (Fig. 4E-F).

**Figure 4.**
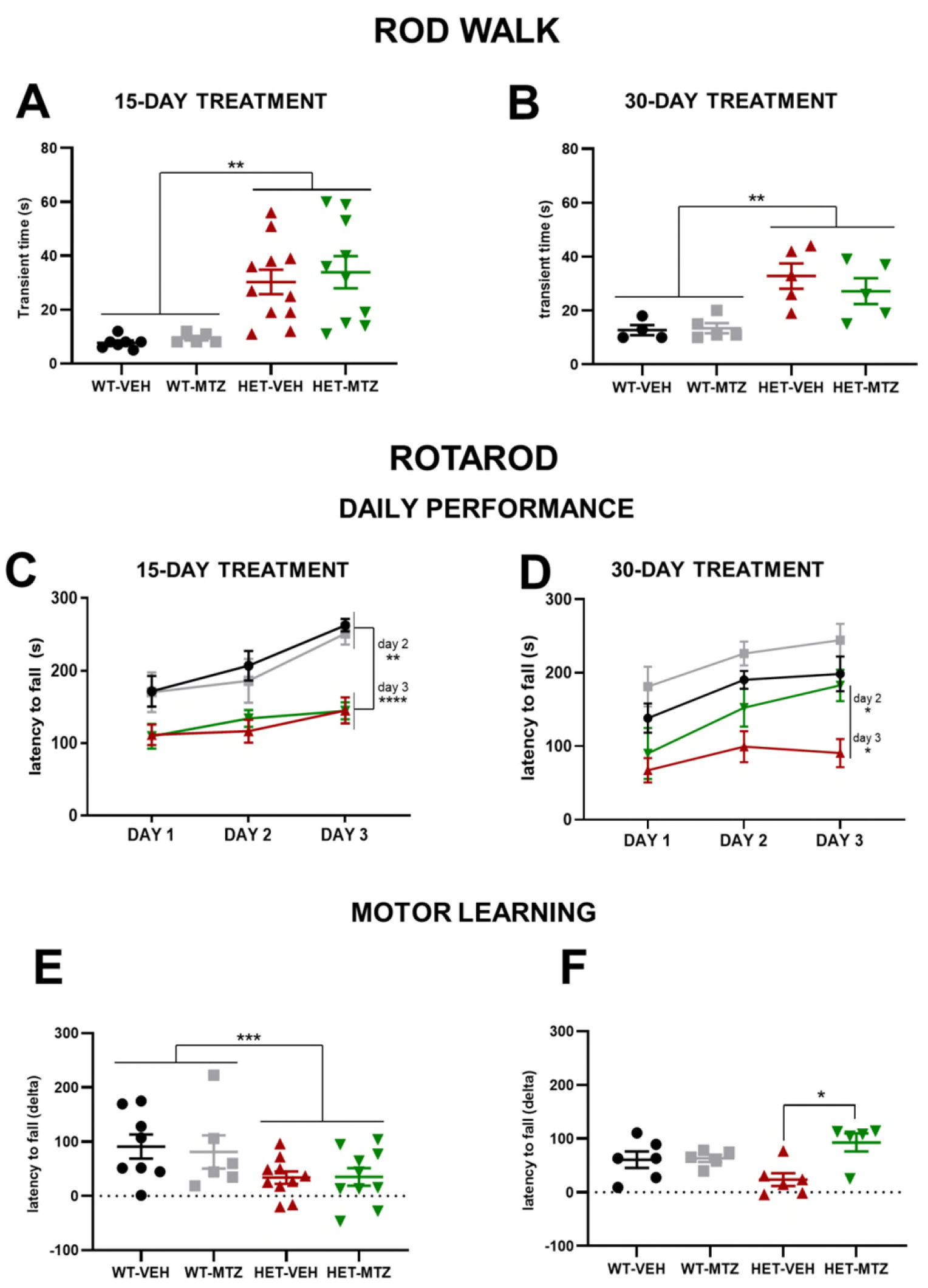
General motor performance of 11-week-old *Mecp2*^tm1.1Bird^ female mice after 15 and 30-day treatment with MTZ. **(A-B)** Time spent in travelling across the 8-mm rod walk is shown. **(C-D)** Daily performance at the Accelerating rotarod is shown. **(E-F)** Motor learning, measured as delta values of latency to fall (third day’s value – second day’s value) at the Accelerating rotarod, is shown. All data are expressed as mean ± SEM and all individual values are showed, n = 8-11 mice/group for 15-day analysis and 5-6 mice/group for 30-day analysis. A standard two-way ANOVA followed by Tukey’s multiple comparisons test was performed. For analysis of daily performance on the Rotarod, we realized a three-way ANOVA followed by Tukey’s multiple comparisons test. ns: p> 0.05 (not indicated), *: p ≤ 0.05, **: p ≤ 0.01, ***: p ≤ 0.001.

### Daily injections affect HET mice but not WT mice

Together with relatively pure motor tests, we aimed to include also a specific somatosensory test with a translational value in the behavioural characterization. To do this, we performed the elevated plus maze (EPM), which we have previously reported to be due to altered whiskers sensitivity in this mouse model *(21)*. Surprisingly, we observed that the phenotype observed in untreated mice (Fig. 1G), consisting in an aberrant preference for the open arms, was no longer present after treatments for both 15 and 30 days (Fig. 5A-B). Not only, but the behaviour of HET mice was exactly the opposite, as they spent less time than WT mice exploring the open arms (even if the difference was statistically significant only after 15 days). A comparison between untreated and untreated WT and HET mice revealed an effect of the daily injections of vehicle (VEH) only in HET mice (Fig. 5C). This probably reflects an increase in anxiety levels in HET mice due to the stressful effect of repeated injections, as these mice could be less able to manage stress than WT mice. Unfortunately, these disruptive effects of chronic vehicle injection precluded us to use EPM to evaluate effects of MTZ on HET mice.

**Figure 5.**
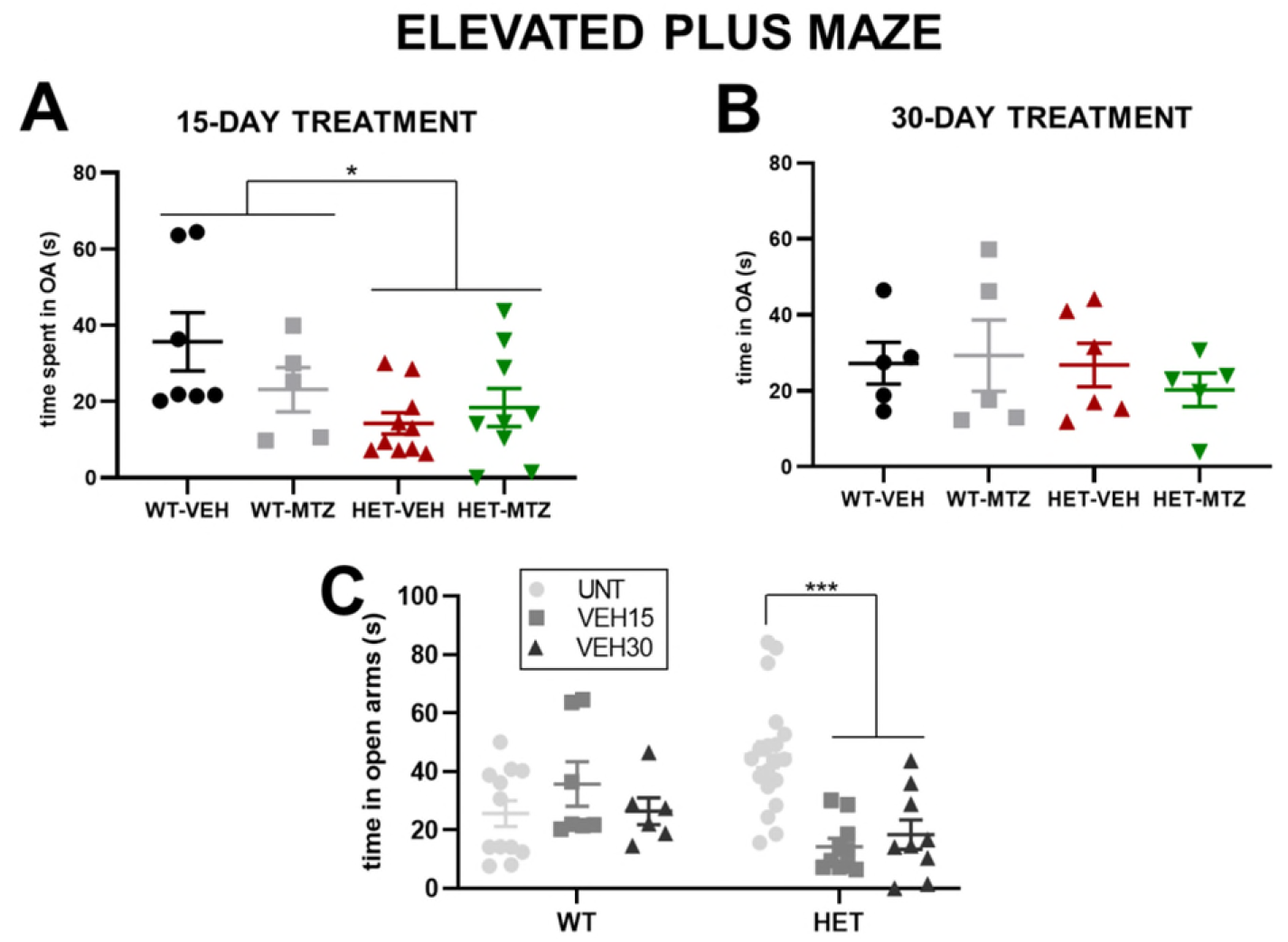
Somatosensory deficits in *Mecp2*^tm1.1Bird^ young female mice. **(A)** Time spent in the open arms of the Elevated plus maze (EPM) after a 15-day treatment with MTZ 10 mg/kg is shown. **(B)** Time spent in the open arms of the EPM after a 30-day treatment with MTZ 10 mg/kg is shown. **(C)** Comparison among untreated (UNT) and treated WT and HET mice treated with vehicle (VEH) which demonstrates and effect of injections only in HET mice. All data are expressed as mean ± SEM and all individual values are showed, n = 8-10 mice/group for 15-day analysis and 5-6 mice/group for 30-day analysis. A standard two-way ANOVA followed by Tukey’s multiple comparisons test was performed. ns: p> 0.05 (not indicated), *: p ≤ 0.05, **: p ≤ 0.01, ***: p ≤ 0.001.

### MTZ normalizes perineuronal nets surrounding PV neurons in the motor cortex

After having performed extensive behavioural testing, we dissected brains of treated WT and HET mice and proceeded with histological analyses. Considering the behavioural results obtained, we decided to focus on the primary motor cortex, which is a main brain area for the motor learning assessed in the accelerating rotarod. Following a previous study in which we had demonstrated that MTZ can normalize some alterations in parvalbuminergic (PV) interneurons *(21)*, we decided to analyse this neuronal type in young HET mice. In addition, we also evaluated the perineuronal nets (PNNs), which are structures of the extracellular matrix often associated with inhibitory neurons such as PV cells, and whose structure is altered in RTT animal models *(39, 40)*. We did not observe any difference in PV cell’s soma area and cell density between WT-VEH and HET-VEH (Fig. 6D-E). We also analysed the interneuronal distance among PV cells, which reflects their distribution within cortical layers, but we did not observe any difference between WT-VEH and HET-VEH (Fig. 6F). We then evaluated the immunoreactivity of PV cells, and observed a significant decrease of immunfluorescence intensity levels in HET-VEH mice compared to WT-VEH at the end of the 15-day treatment with the vehicle (Fig. 6G). Remarkably, this phenotype was completely rescued by treatment with MTZ, as HET-MTZ mice show PV immunoreactivity levels comparable to those of WT-VEH mice (Fig. 6G). Furthermore, we observed that PNNs were thicker in HET-VEH mice, and that MTZ normalized PNNs thickness (Fig. 6H). Regarding the 30-day treatment, we did not observe the phenotype of reduced PV immunoreactivity levels, but we did confirm the PNN-related phenotype and its complete rescue upon MTZ treatment (Fig. 6I-J).

**Figure 6.**
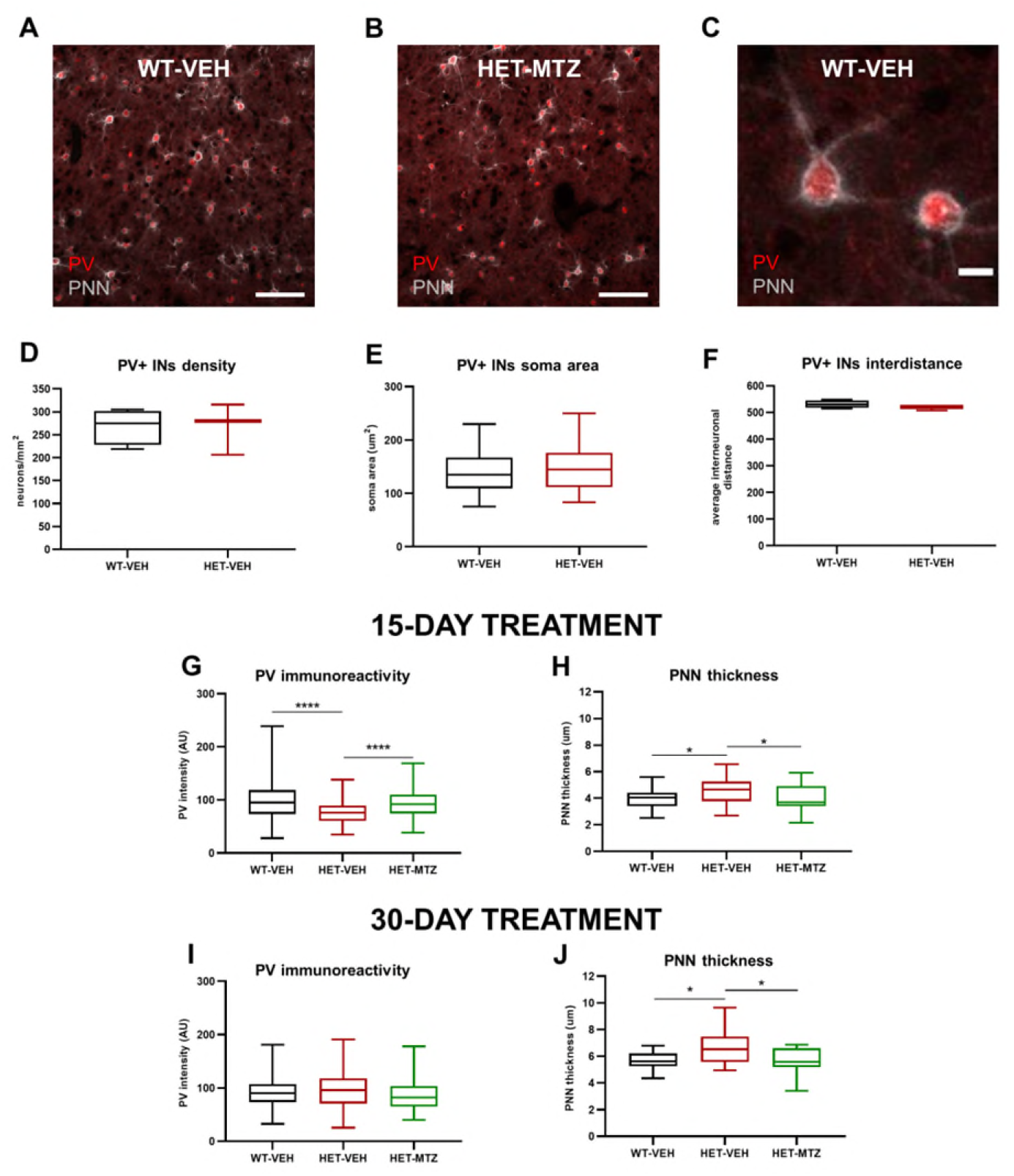
Analysis of parvalbumin-positive interneurons (PV+ INs) and perineuronal nets (PNNs) in the primary motor cortex of 13-week-old *Mecp2*^tm1.1Bird^ female mice after treatment with MTZ (10 mg/kg) for 15 or 30 days. **(A-B)** Examples of PV and PNN staining in a primary motor cortex section (scale bar = 100 μm) of a WT mice treated with VEH and a HET mouse treated with MTZ. **(C)** Detail of two PV+ interneurons (INs) associated to PNNs in a primary motor cortex section (scale bar = 10 μm) of a WT-VEH mouse. **(D)** Density of PV+ INs in the 1^ary^ motor cortex of mice treated for 15 days. N = 10 slices/mouse and 3-4 mice/group. **(E)** Soma area of PV INs in the 1^ary^ motor cortex of mice treated for 15 days. N = 10 neurons/mouse and 4 mice/group. **(F)** Average distance among PV+ interneurons in the 1^ary^ motor cortex of mice treated for 15 days. N = 2-3 slices/mouse and 3-4 mice/group. **(G)** Immunoreactivity of PV INs’ somata in the 1^ary^ motor cortex of mice treated for 15 days. N = 10 neurons/mouse and 3-4 mice/group. **(H)** Average thickness of PNNs surrounding PV+ INs in the 1^ary^ motor cortex of mice treated for 15 days. N = 10 neurons/mouse and 4 mice/group. **(I)** Immunoreactivity of PV INs’ somata in the 1^ary^ motor cortex of mice treated for 30 days. N = 10 neurons/mouse and 3-4 mice/group. **(J)** Average thickness of PNNs surrounding PV+ INs in the 1^ary^ motor cortex of mice treated for 30 days. N = 10 neurons/mouse and 4 mice/group. All data are represented as median ± interquartile range and maximum and minimum data points. According to results of Shapiro-Wilk test, either one-way ANOVA, followed by Tukey’s post-hoc test, or Kruskal-Wallis test, followed by Dunn’s post-hoc test, was performed. Multiple selected comparisons comprehended: WT-VEH vs WT-MTZ, WT-VEH vs HET-VEH and HET-VEH vs HET-MTZ. ns: p> 0.05 (not indicated), *: p ≤ 0.05, **: p ≤ 0.01, ***: p ≤ 0.001.

## Discussion

In this study, we carried out systematic behavioural characterization of the sensorimotor developmental maturation, with an additional short-memory task, of HET *Mecp2*^tm1.1Bird^ female mice at two different ages (6 and 11 weeks) and found previously unknown defective phenotypes. In particular, we found significant RTT-like signs of developmental disruption already at 6 weeks of age, such as increased body weight, hindlimb clasping, impaired general motor coordination (as showed at the rod walk test) and aberrant preference for open arms in the elevated plus maze at both ages. In addition, we also detected significant motor impairments in the accelerating rotarod and the adhesive patch removal tests, as well as in the short-term memory while performing the four-different objects task (4-DOT). The progressive age-dependent worsening of the behavioural phenotype from basic sensorimotor deficit, neuromotor coordination and skilled learning suggests that cortico-subcortical regions are progressively comprised in female HET mice before reaching adulthood and recapitulates the complexity of the neurological phenotype observed in RTT children.

Results of this analysis indicated that 11 weeks was a better age for assessing the effects of the drugs on the sensorimotor development and therefore, we started treatment with MTZ at 9 weeks and compared two different treatment durations, namely 15 and 30 days, at the daily dose of 10 mg/kg (the equivalent to the maximum dose admitted in humans). We found that the longer MTZ treatment showed higher efficacy than the shorter one. Indeed, 30 days, but not 15 days of treatment with MTZ induced remission of body weight increase. Interestingly, 30-day MTZ treatment did not rescue the basal motor coordination deficit in the rod task, but it was effective in recovering both hindlimb clasping and motor learning deficits as evaluated in the rotarod task. Motor learning in the accelerating rotarod test requires functional activation of cortical-subcortical inputs and therefore we decided to investigate maturation of inhibitory neurons in the primary motor cortex, which have previously shown to show alterations in HET *Mecp2*^tm1.1Bird^ mice *(21)*. As the alterations in perineuronal nets (PNNs) in the primary motor cortex were fully recovered by a 30-day long MTZ treatment, we propose correction of the developmental programme for interneuron maturation in the motor cortex as one underlying mechanism for the recovery of motor signs induced by MTZ. Taken together, the results obtained here clearly show that MTZ treatment for at least 30 days within a time window corresponding to late adolescence was efficacious, and strongly support MTZ as a candidate to treat young girls affected by RTT in a future clinical trial.

### 1. Detection of novel behavioural phenotypes at 6 and 11 weeks

As relatively few information on young *Mecp2*^tm1.1Bird^ heterozygous (HET) female mice phenotype were previously available, we decided to perform an extensive behavioural characterization at 6 and 11 weeks of age, which correspond to “early adolescence” and “late adolescence/young adulthood”, respectively *(28)*. The behavioural characterization of HET mice at 6 and 11 weeks of age revealed several previously unknown alterations in the sensorimotor domain, revealing these mice as a good model to test drugs also at young ages, when plasticity processes are more active.

We found that body weight was significantly increased in HET mice as soon as 6 weeks of age, which confirmed previous observations by Vogel Ciernia et al. in HET mice at 6 and 11 weeks *(26)*. Hindlimb clasping was present in most HET mice at both 6 and 11 weeks. This very typical progressive RTT-like sign, which resembles the hand clasping found in patients, has previously been documented by several laboratories in both male *Mecp2*^y/-^ *(20, 22, 41, 42)* and female *Mecp2*^*+/-*^ murine models *(21, 26, 39)*, but in the latter model, the earliest age at which it had been observed so far was the 8^th^ postnatal week *(26)*. Hence, our findings anticipate the onset of the hindlimb clasping by two weeks, i.e., to 6 weeks of age, with respect to previous reports.

Regarding other motor phenotypes, we observed that the phenotype described in adult HET mice at the Horizontal bars test *(21)* is not present in 11-month-old HET mice, while both accelerating rotarod and rod walk revealed consistent deficits at 6 and 11 weeks of age, like previously reported in adults *(21)*. Regarding the accelerating rotarod, we specifically showed that HET mice present impairment not only of daily performance, but also of motor learning within the three consecutive days of test. The deficits we observed in the accelerating rotarod performance are in accordance with previous studies reporting that this impairment is present in female HET mice as early as 5 weeks from birth *(43, 44)*. In addition, at 11 weeks of age, we observed a previously undescribed phenotype in the adhesive patch removal task, which evaluates the fine motricity of fore limbs.

We further show that the aberrant preference for the open arms commonly observed in both male and female adult mice when tested at the EPM *(20, 22, 26, 41, 45)* is already manifest in young mice at both 6 and 11 weeks of age. In a previous study, we demonstrated that this behaviour likely implicates sensory hypersensitivity at the whiskers, as it was completely abolished by clipping mouse whiskers *(21)*. Finally, we tested mice at the four-different object task (4-DOT), as also the cognitive domain has been poorly studied in RTT mouse models. The way in which this test was performed allows the evaluation of short-term memory of mice associated with tactile and visual stimuli *(31)*. We first used the 4-DOT in untreated mice at both 6 and 11 weeks of age and showed a cognitive deficit only in older mice.

According to results of this behavioural characterization, we decided to exclude both patch adhesive removal test and the 4-DOT for the battery of tests to perform after the pharmacological treatment, as we wanted to test the rescuing effect of the drug on already established defects. In addition, we aimed to use a low number of tests after the pharmacological treatment to limit the stress on mice, considering that i.p. injections were performed daily and hence represented already a stressing continuous stimulus.

### 2. Physiological significance of the effects of MTZ in young HET mice

Nowadays, pharmacological approaches together with physical therapies are the only strategy to improve the quality of life of RTT patients, through the recovery or alleviation of symptomatology. However, just a few drugs have demonstrated this potential so far, and none of them has arrived to reach the clinical consensus level to treat RTT as a whole. We chose MTZ as a candidate because of some of its pharmacological and clinical features. Above all, this antidepressant satisfies the most important requirement for any drug candidate to treat RTT: MTZ is a relatively safe drug. Since its approval for the treatment of major depression in 1996, it has shown an excellent safety profile *(17)*. Moreover, a recent study comparing twenty-one well-known antidepressants has concluded that MTZ is one of the most suitable ones to treat major depression disorders in terms of tolerability *(16)*. Finally, its numerous off-label uses *(46)* are also evidence of the few side effects exerted by MTZ. Most importantly, we previously provide evidence of the high tolerability to MTZ in 40 adult RTT patients chronically treated for several years (up to 5 years) *(21)*. In the current study, we have largely confirmed MTZ safety. In fact, we have observed no adverse effects in young female WT-MTZ and HET-MTZ mice, as they did not show any phenotypic worsening compared to VEH controls, even when treated for 30 days with a high dose of MTZ (10 mg/kg, the equivalent to the maximum dose admitted in humans, i.e., 50 mg/day). These findings have important implications for future research, as it will be possible to test a reasonably wide range of dosages in a randomized clinical trial in young RTT patients.

Concerning the possible mechanisms of action of MTZ, it should be noted that this antidepressant has a unique profile. It is characterized by a relatively rapid onset of action, high response and remission rates, a favourable side-effect profile, and several unique therapeutic benefits over other antidepressants *(47)*. So far, 18 receptors have been identified as targets of MTZ in the brain *(15)*, but the effects exerted on the nervous system following their activation by MTZ are just partially known. MTZ shows its highest affinity for histaminergic H1 receptor, which is responsible for sedation when used at low doses *(48)*. Regarding monoaminergic systems, which are generally downregulated in RTT patients and mouse models *(25, 49, 50)*, MTZ enhances 5-HT and NA transmission through different mechanisms. At the presynaptic level, it antagonizes α-2 auto- and hetero-adrenoreceptors, leading to an increase in NA release, while at the postsynaptic level it antagonizes 5-HT_2_ and 5-HT_3_ receptors, promoting an increase in the release of serotonin leading to an indirect enhancement of 5-HT_1A_-mediated serotonergic transmission. In addition, NA released in the raphe nuclei further stimulates postsynaptic α-1 receptors, causing 5-HT release from downstream axon terminals such as those in the cortex *(51, 52)*. Specific agonists of the 5-HT_1A_ receptor have shown promising abilities to alleviate brainstem and extrapyramidal dysfunction in preclinical studies on RTT autonomic phenotypes *(23)*. However, some of the most prominent among these drugs have also been shown to antagonize partially or totally DA receptors. This makes them not suitable for RTT, as also the dopaminergic system is generally altered in patients and mouse models. For example, low spinal fluid DA levels, which are assumed to be a marker for central DA levels, have been found in women with known *MECP2* mutations and meeting the clinical criteria for RTT *(25)*. Furthermore, the presence of a DA deficit is further supported by the fact that, later in life, RTT patients develop Parkinsonian features *(12)*. Regarding these aspects, MTZ is an optimal candidate for RTT, as it not only enhances 5-HT transmission through 1A receptors (with minimal occurrence of serotonergic side effects) but it also increases DA transmission *(51, 53)*. On the other side, it is known that 5-HT regulates motor networks facilitating motor skill, motor cortex plasticity and motor output *(54)* and it has been recently found that 5-HT has a major role in pharmacological recovery of motor performance at the accelerating rotarod *(44)*. Moreover, selective 5-HT reuptake inhibitors (SSRIs) has been demonstrated to enhance motor skill learning and plasticity *(55–57)*. It is therefore conceivable that the rescue observed in the motor domain after MTZ treatment in both HET mice and RTT treatments may mainly involve the enhancement of 5-HT transmission, but also the normalization of DA levels in the brain, along with the rescue of the inhibitory interneurons function. In fact, another mechanism investigated in this study is related to the maturation of parvalbuminergic (PV) neuronal networks. PV cells are interneurons playing a main role in GABAergic inhibition and having been associated with RTT symptomatology and related phenotypes in mice *(40, 58, 59)*. In a previous histological analysis on MTZ-treated adult *Mecp2*^tm1.1Bird^ female mice, we found alterations in PV expression levels and increased PV-positive cell density in several brain areas, comprising motor, somatosensory cortex and amygdala, some of which were normalized by MTZ *(21)*. Here, while we detected also in young HET mice a significant reduction in the PV immunoreactivity in the primary motor cortex, we did not observe any alterations in the PV-positive cell density, soma size or intercellular distance. Considering the progressive nature of RTT, it is possible that, besides the reduction in PV expression levels, other alterations on parvalbumin networks may not become evident until adult ages.

Together with PV networks, perineuronal nets (PNNs), which surround mainly PV interneurons in the brain cortex, have been also associated with RTT physiopathology *(21, 40, 59)*). In this case, the hypothesis to verify was that an abnormal early development of PNNs, which physiologically limit synaptic activity, could contribute to behavioural RTT-like phenotypes. In our analysis we did not find differences neither in the PNNs density, nor in their immunoreactivity (*data not shown*) in 11-week-old *Mecp2*^tm1.1Bird^ female mice. However, we found that PNN thickness was significantly increased in HET control mice and could be corrected by a 30-day MTZ treatment. The beneficial effect of MTZ to promote PV expression and normalization of PV cells engulfment by perineuronal nets in the motor cortex is in agreement with our previous studies in adult female HET mice and provides a strong indication of one possible mechanism of action of this drug for the correction of aberrant RTT-like motor behaviours.

### 3. Limitations of the study

Besides the encouraging results described above, this study also presents some limitations. First, considering that RTT is a progressive disorder, it would be desirable to treat RTT patients as early as possible. However, the hardly measurable phenotype at 6 weeks, as compared to 11 weeks, prevented us to test the efficacy of MTZ at the earlier stage investigated in this study. Second, this study highlighted some limitations in the effects of MTZ, as it was not able to rescue deficits observed at the rod walk, which evaluates the general motor coordination. At present, it remains open the possibility that this lack of efficacy of MTZ treatment may be due either to unresponsiveness to the drug of the underlying neuronal circuitries, which for the rod walk include also the cerebellum, or to the fact that longer treatment times may be required to show any improvement in these deficits. As one additional limitation, we observed that potential effects of MTZ in the Elevated plus maze were covered but stress effects of daily i.p. injections.

### 4. Conclusions, a lesson for a future clinical trial in RTT children and adolescents

From this study, MTZ emerges as a drug which is well tolerated also in juvenile mouse models of RTT, and with strong potential for long-term chronic treatment of children and adolescent with RTT. In fact, the combination of the results presented here and our recently published investigation on adult female mice and patients *(21)* clearly indicates that the time window for a successful outcome of the treatment of RTT with MTZ must be large enough (months to years) to allow improvement of the pathological deficits in the neural circuits underlying RTT and, therefore, also in quality of life of patients.

## Acknowledgements

We thank Olga Carofiglio for her collaboration in the immunohistological analyses. We thank ProRett Ricerca ONLUS for the generous support of this study. JFG was a recipient of a full PhD bursary of the University of Trieste, Doctorate School in Molecular Biomedicine.

## FIGURES

**Suppl. Fig. 1.**
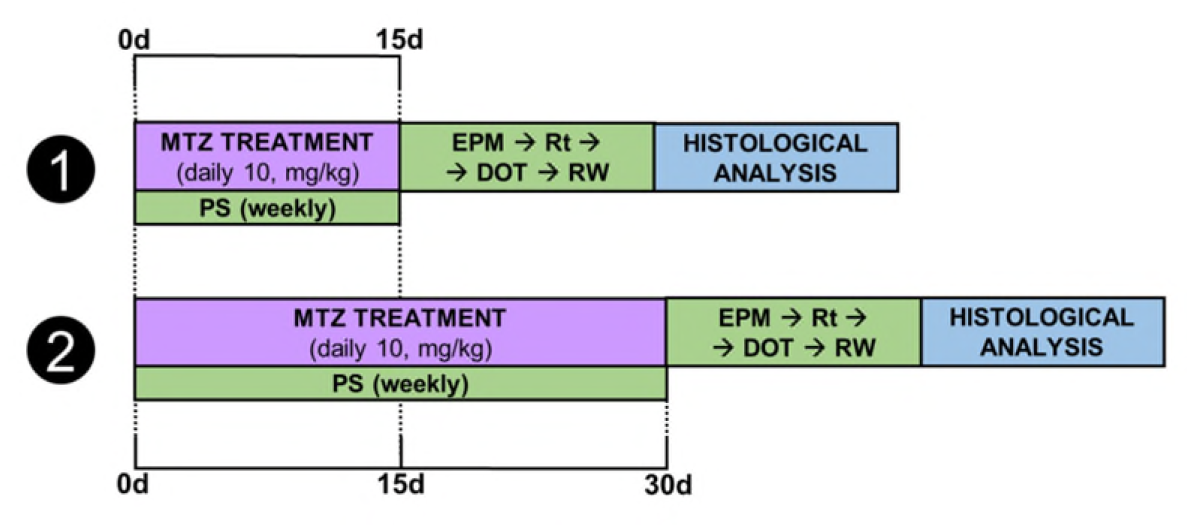
Timelines of experimental designs. The two different general protocols are shown. Treatment with mirtazapine (MTZ) or vehicle (VEH) was initiated when *Mecp2*^tm1.1Bird^ heterozygous (HET) female mice were 9 weeks old. In both cases, general health was evaluated weekly (beginning at the first day of treatment) through a phenotypic scoring (PS) modified from Guy et al. (2007). MTZ was injected intraperitoneally (i.p.) every day and dosage administered in each single injection was always the same (10 mg/kg). After both 15-day and 30-day treatments, behavioural tests were performed following the same order (EPM: Elevated-plus maze; Rt: Accelerating rotarod; DOT: 4-different object task; RW: Rod walk test). Beginning from the day after the last injection, one test was performed per day to avoid excessive stress and interactions. In all cases, mice brains were dissected and prepared for different histological analysis at the end of post-treatment behavioural testing (see Materials and Methods’ specific section).

**Suppl. Fig. 2.**
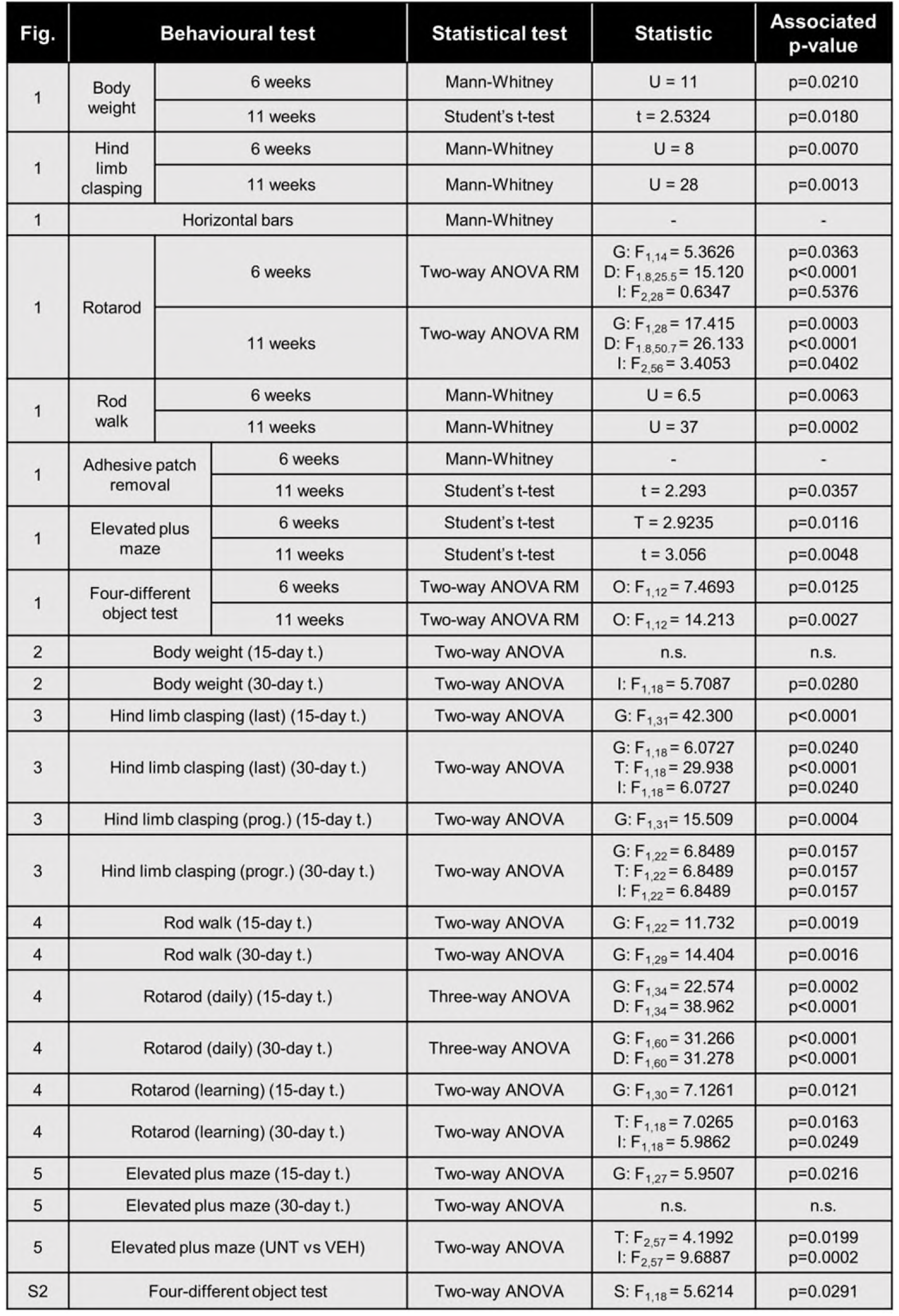
Details of statistical analyses. For each behavioural test, it is indicated the type of statistical test performed, the value of the statistic (U, t or F) and the associated p-value. These parameters are shown only for statistically significant results, while n.s. is used to indicate that no significant results were found in that specific behavioural test. For two-way ANOVA analyses, it is also indicated the factor to which the statistic is associated: genotype (G), day (D), object (O), treatment (T) or segment (S). In addition, statistical parameters of interaction of factors (I) of two-way ANOVA are also indicated. For p-values associated to specific multiple comparisons, see the correspondent figure.

## REFERENCES

1. A. Rett, On a remarkable syndrome of cerebral atrophy associated with hyperammonaemia in childhood, Wien. Med. Wochenschr. 166, 322–324 (2016).

2. B. Hagberg, J. Aicardi, K. Dias, O. Ramos, A progressive syndrome of autism, dementia, ataxia, and loss of purposeful hand use in girls: Rett’s syndrome: Report of 35 cases, Ann. Neurol. 14, 471–479 (1983).

3. S. Fehr, A. Bebbington, N. Nassar, J. Downs, G. M. Ronen, N. De Klerk, H. Leonard, Trends in the diagnosis of Rett syndrome in Australia, Pediatr. Res. 70, 313–319 (2011).

4. R. E. Amir, I. B. Van den Veyver, M. Wan, C. Q. Tran, U. Francke, H. Y. Zoghbi, Rett syndrome is caused by mutations in X-linked MECP2, encoding methyl-CpG-binding protein 2, Nat. Genet. 23, 185–188 (1999).

5. Neul, W. E. Kaufmann, D. G. Glaze, J. Christodoulou, A. J. Clarke, N. Bahi-Buisson, H. Leonard, M. E. S. Bailey, N. C. Schanen, M. Zappella, A. Renieri, P. Huppke, A. K. Percy, for the RettSearch Consortium (Members listed in the Appendix), Rett syndrome: revised diagnostic criteria and nomenclature, Ann. Neurol. 68, 944–950 (2010).

6. V. A. Cuddapah, R. B. Pillai, K. V. Shekar, J. B. Lane, K. J. Motil, S. A. Skinner, D. C. Tarquinio, D. G. Glaze, G. McGwin, W. E. Kaufmann, A. K. Percy, J. L. Neul, M. L. Olsen, Methyl-CpG-binding protein 2 (MECP2) mutation type is associated with disease severity in Rett syndrome, J. Med. Genet. 51, 152–158 (2014).

7. A. Erlandson, B. Hagberg, MECP2 abnormality phenotypes: clinicopathologic area with broad variability, J. Child Neurol. 20, 727–732 (2005).

8. D. R. Connolly, Z. Zhou, Genomic insights into MeCP2 function: A role for the maintenance of chromatin architecture, Curr. Opin. Neurobiol. 59, 174–179 (2019).

9. F. Della Ragione, M. Vacca, S. Fioriniello, G. Pepe, M. D’Esposito, MECP2, a multi-talented modulator of chromatin architecture, Brief. Funct. Genomics, elw023 (2016).

10. M. Chahrour, H. Y. Zoghbi, The story of Rett Syndrome: from clinic to neurobiology, Neuron 56, 422–437 (2007).

11. L. Jian, L. Nagarajan, N. de Klerk, D. Ravine, C. Bower, A. Anderson, S. Williamson, J. Christodoulou, H. Leonard, Predictors of seizure onset in Rett syndrome, J. Pediatr. 149, 542-547.e3 (2006).

12. E. Roze, V. Cochen, S. Sangla, T. Bienvenu, A. Roubergue, S. Leu-Semenescu, M. Vidaihet, Rett syndrome: an overlooked diagnosis in women with stereotypic hand movements, psychomotor retardation, Parkinsonism, and dystonia?, Mov. Disord. Off. J. Mov. Disord. Soc. 22, 387–389 (2007).

13. A. K. Percy, Progress in Rett Syndrome: from discovery to clinical trials, Wien. Med. Wochenschr. 166, 325–332 (2016).

14. H. Zoghbi, A. Percy, D. Glaze, I. Butler, V. Riccardi, Reduction of biogenic amine leveles in the Rett syndrome, (1985).

15. S. A. K. Anttila, E. V. J. Leinonen, A Review of the Pharmacological and Clinical Profile of Mirtazapine, CNS Drug Rev. 7, 249–264 (2006).

16. A. Cipriani, T. A. Furukawa, G. Salanti, A. Chaimani, L. Z. Atkinson, Y. Ogawa, S. Leucht, H. G. Ruhe, E. H. Turner, J. P. T. Higgins, M. Egger, N. Takeshima, Y. Hayasaka, H. Imai, K. Shinohara, A. Tajika, J. P. A. Ioannidis, J. R. Geddes, Comparative efficacy and acceptability of 21 antidepressant drugs for the acute treatment of adults with major depressive disorder: a systematic review and network meta-analysis, The Lancet 391, 1357–1366 (2018).

17. A. Szegedi, N. Schwertfeger, Mirtazapine: a review of its clinical efficacy and tolerability, Expert Opin. Pharmacother. 6, 631–641 (2005).

18. G. D. Burrows, C. M. E. Kremer, Mirtazapine: clinical advantages in the treatment of depression, J. Clin. Psychopharmacol. 17, 34S–39S (1997).

19. Hartmann, Peter, Mirtazapine: a newer antidepressant, Am. Fam. Physician 59, 159–161 (1999).

20. T. Bittolo, C. A. Raminelli, C. Deiana, G. Baj, V. Vaghi, S. Ferrazzo, A. Bernareggi, E. Tongiorgi, Pharmacological treatment with mirtazapine rescues cortical atrophy and respiratory deficits in MeCP2 null mice, Sci. Rep. 6, 19796 (2016).

21. J. Flores Gutiérrez, C. De Felice, G. Natali, S. Leoncini, C. Signorini, J. Hayek, E. Tongiorgi, Protective role of mirtazapine in adult female Mecp2+/-mice and patients with Rett syndrome, J. Neurodev. Disord. 12, 26 (2020).

22. J. Guy, B. Hendrich, M. Holmes, J. E. Martin, A. Bird, A mouse Mecp2-null mutation causes neurological symptoms that mimic Rett syndrome, Nat. Genet. 27, 322–326 (2001).

23. A. P. Abdala, D. T. Lioy, S. K. Garg, S. J. Knopp, J. F. R. Paton, J. M. Bissonnette, Effect of Sarizotan, a 5-HT _1a_ and D2-Like Receptor Agonist, on Respiration in Three Mouse Models of Rett Syndrome, Am. J. Respir. Cell Mol. Biol. 50, 1031–1039 (2014).

24. M. Horiuchi, L. Smith, I. Maezawa, L.-W. Jin, CX3CR1 ablation ameliorates motor and respiratory dysfunctions and improves survival of a Rett syndrome mouse model, Brain. Behav. Immun. 60, 106–116 (2017).

25. R. C. Samaco, C. Mandel-Brehm, H.-T. Chao, C. S. Ward, S. L. Fyffe-Maricich, J. Ren, K. Hyland, C. Thaller, S. M. Maricich, P. Humphreys, J. J. Greer, A. Percy, D. G. Glaze, H. Y. Zoghbi, J. L. Neul, Loss of MeCP2 in aminergic neurons causes cell-autonomous defects in neurotransmitter synthesis and specific behavioral abnormalities, Proc. Natl. Acad. Sci. 106, 21966–21971 (2009).

26. A. Vogel Ciernia, M. C. Pride, B. Durbin-Johnson, A. Noronha, A. Chang, D. H. Yasui, J. N. Crawley, J. M. LaSalle, Early motor phenotype detection in a female mouse model of Rett syndrome is improved by cross-fostering, Hum. Mol. Genet. 26, 1839–1854 (2017).

27. J. Guy, J. Gan, J. Selfridge, S. Cobb, A. Bird, Reversal of neurological defects in a mouse model of Rett syndrome, Science 315, 1143–1147 (2007).

28. S. Dutta, P. Sengupta, Men and mice: Relating their ages, Life Sci. 152, 244–248 (2016).

29. S. C. Landis, S. G. Amara, K. Asadullah, C. P. Austin, R. Blumenstein, E. W. Bradley, R. G. Crystal, R. B. Darnell, R. J. Ferrante, H. Fillit, R. Finkelstein, M. Fisher, H. E. Gendelman, R. M. Golub, J. L. Goudreau, R. A. Gross, A. K. Gubitz, S. E. Hesterlee, D. W. Howells, J. Huguenard, K. Kelner, W. Koroshetz, D. Krainc, S. E. Lazic, M. S. Levine, M. R. Macleod, J. M. McCall, R. T. Moxley III, K. Narasimhan, L. J. Noble, S. Perrin, J. D. Porter, O. Steward, E. Unger, U. Utz, S. D. Silberberg, A call for transparent reporting to optimize the predictive value of preclinical research, Nature 490, 187–191 (2012).

30. A. Nair, S. Jacob, A simple practice guide for dose conversion between animals and human, J. Basic Clin. Pharm. 7, 27 (2016).

31. S. Sannino, F. Russo, G. Torromino, V. Pendolino, P. Calabresi, E. De Leonibus, Role of the dorsal hippocampus in object memory load, Learn. Mem. 19, 211–218 (2012).

32. R. M. J. Deacon, Measuring Motor Coordination in Mice, J. Vis. Exp., 2609 (2013).

33. V. Bouet, M. Boulouard, J. Toutain, D. Divoux, M. Bernaudin, P. Schumann-Bard, T. Freret, The adhesive removal test: a sensitive method to assess sensorimotor deficits in mice, Nat. Protoc. 4, 1560–1564 (2009).

34. G. Paxinos, K. Franklin, The Mouse Brain in Stereotaxic Coordinates, Compact (Academic Press, 3rd Edition., 2008).

35. J. Schindelin, I. Arganda-Carreras, E. Frise, V. Kaynig, M. Longair, T. Pietzsch, S. Preibisch, C. Rueden, S. Saalfeld, B. Schmid, J.-Y. Tinevez, D. J. White, V. Hartenstein, K. Eliceiri, P. Tomancak, A. Cardona, Fiji: an open-source platform for biological-image analysis, Nat. Methods 9, 676–682 (2012).

36. D. M. Katz, J. E. Berger-Sweeney, J. H. Eubanks, M. J. Justice, J. L. Neul, L. Pozzo-Miller, M. E. Blue, D. Christian, J. N. Crawley, M. Giustetto, J. Guy, C. J. Howell, M. Kron, S. B. Nelson, R. C. Samaco, L. R. Schaevitz, C. St. Hillaire-Clarke, J. L. Young, H. Y. Zoghbi, L. A. Mamounas, Preclinical research in Rett syndrome: setting the foundation for translational success, Dis. Model. Mech. 5, 733–745 (2012).

37. R. C. Samaco, C. M. McGraw, C. S. Ward, Y. Sun, J. L. Neul, H. Y. Zoghbi, Female Mecp2+/-mice display robust behavioral deficits on two different genetic backgrounds providing a framework for pre-clinical studies, Hum. Mol. Genet. 22, 96–109 (2013).

38. G. S. Tomassy, E. De Leonibus, D. Jabaudon, S. Lodato, C. Alfano, A. Mele, J. D. Macklis, M. Studer, Area-specific temporal control of corticospinal motor neuron differentiation by COUP-TFI, Proc. Natl. Acad. Sci. 107, 3576–3581 (2010).

39. A. Patrizi, N. Picard, A. J. Simon, G. Gunner, E. Centofante, N. A. Andrews, M. Fagiolini, Chronic Administration of the N-Methyl-D-Aspartate Receptor Antagonist Ketamine Improves Rett Syndrome Phenotype, Biol. Psychiatry 79, 755–764 (2016).

40. A. Patrizi, P. N. Awad, B. Chattopadhyaya, C. Li, G. Di Cristo, M. Fagiolini, Accelerated Hyper-Maturation of Parvalbumin Circuits in the Absence of MeCP2, Cereb. Cortex 30, 256–268 (2020).

41. C. Cobolli Gigli, L. Scaramuzza, A. Gandaglia, E. Bellini, M. Gabaglio, D. Parolaro, C. Kilstrup-Nielsen, N. Landsberger, F. Bedogni, M. D’Esposito, Ed. MeCP2 Related Studies Benefit from the Use of CD1 as Genetic Background, PLOS ONE 11, e0153473 (2016).

42. G. J. Pelka, C. M. Watson, T. Radziewic, M. Hayward, H. Lahooti, J. Christodoulou, P. P. L. Tam, Mecp2 deficiency is associated with learning and cognitive deficits and altered gene activity in the hippocampal region of mice, Brain 129, 887–898 (2006).

43. M. Santos, A. Silva-Fernandes, P. Oliveira, N. Sousa, P. Maciel, Evidence for abnormal early development in a mouse model of Rett syndrome, Genes Brain Behav. 6, 277–286 (2007).

44. C. Villani, G. Sacchetti, R. Bagnati, A. Passoni, F. Fusco, M. Carli, R. W. Invernizzi, Lovastatin fails to improve motor performance and survival in methyl-CpG-binding protein2-null mice,, 14 (2016).

45. C. S. Ward, E. M. Arvide, T.-W. Huang, J. Yoo, J. L. Noebels, J. L. Neul, MeCP2 Is Critical within HoxB1-Derived Tissues of Mice for Normal Lifespan, J. Neurosci. 31, 10359–10370 (2011).

46. T. N. Jilani, A. Saadabadi, in StatPearls, (StatPearls Publishing, Treasure Island (FL), 2019).

47. A. Alam, Z. Voronovich, J. A. Carley, A Review of Therapeutic Uses of Mirtazapine in Psychiatric and Medical Conditions, Prim. Care Companion CNS Disord. 15 (2013), doi:10.4088/PCC.13r01525.

48. J. Fawcett, R. L. Barkin, Review of the results from clinical studies on the efficacy, safety and tolerability of mirtazapine for the treatment of patients with major depression, J. Affect. Disord. 51, 267–285 (1998).

49. M. Santos, T. Summavielle, A. Teixeira-Castro, A. Silva-Fernandes, S. Duarte-Silva, F. Marques, L. Martins, M. Dierssen, P. Oliveira, N. Sousa, P. Maciel, Monoamine deficits in the brain of methyl-CpG binding protein 2 null mice suggest the involvement of the cerebral cortex in early stages of Rett syndrome, Neuroscience 170, 453–467 (2010).

50. T. Temudo, M. Rios, C. Prior, I. Carrilho, M. Santos, P. Maciel, J. Sequeiros, M. Fonseca, J. Monteiro, P. Cabral, J. P. Vieira, A. Ormazabal, R. Artuch, Evaluation of CSF neurotransmitters and folate in 25 patients with Rett disorder and effects of treatment, Brain Dev. 31, 46–51 (2009).

51. K. Nakayama, T. Sakurai, H. Katsu, Mirtazapine increases dopamine release in prefrontal cortex by 5-HT1A receptor activation, Brain Res. Bull. 63, 237–241 (2004).

52. Sthal, Stephen M., Stahl’s Essential Psychopharmacology: Neuroscientific Basis and Practical Applications, Mens Sana Monogr. 8, 146–150 (2010).

53. M. Masana, A. Castañé, N. Santana, A. Bortolozzi, F. Artigas, Noradrenergic antidepressants increase cortical dopamine: Potential use in augmentation strategies, Neuropharmacology 63, 675–684 (2012).

54. C. Vitrac, M. Benoit-Marand, Monoaminergic Modulation of Motor Cortex Function, Front. Neural Circuits 11, 72 (2017).

55. G. Batsikadze, W. Paulus, M.-F. Kuo, M. A. Nitsche, Effect of Serotonin on Paired Associative Stimulation-Induced Plasticity in the Human Motor Cortex, Neuropsychopharmacology 38, 2260–2267 (2013).

56. A. Gerdelat-Mas, I. Loubinoux, D. Tombari, O. Rascol, F. Chollet, M. Simonetta-Moreau, Chronic administration of selective serotonin reuptake inhibitor (SSRI) paroxetine modulates human motor cortex excitability in healthy subjects, NeuroImage 27, 314–322 (2005).

57. I. Loubinoux, D. Tombari, J. Pariente, A. Gerdelat-Mas, X. Franceries, E. Cassol, O. Rascol, J. Pastor, F. Chollet, Modulation of behavior and cortical motor activity in healthy subjects by a chronic administration of a serotonin enhancer, NeuroImage 27, 299–313 (2005).

58. A. Ito-Ishida, K. Ure, H. Chen, J. W. Swann, H. Y. Zoghbi, Loss of MeCP2 in Parvalbumin-and Somatostatin-Expressing Neurons in Mice Leads to Distinct Rett Syndrome-like Phenotypes, Neuron 88, 651–658 (2015).

59. N. Morello, R. Schina, F. Pilotto, M. Phillips, R. Melani, O. Plicato, T. Pizzorusso, L. Pozzo-Miller, M. Giustetto, Loss of Mecp2 causes atypical synaptic and molecular plasticity of parvalbumin-expressing interneurons reflecting Rett syndrome–like sensorimotor defects, eneuro 5, ENEURO.0086-18.2018 (2018).

